# Caffeine-dependent changes of sleep-wake regulation: evidence for adaptation after repeated intake

**DOI:** 10.1101/641480

**Authors:** Janine Weibel, Yu-Shiuan Lin, Hans-Peter Landolt, Corrado Garbazza, Vitaliy Kolodyazhniy, Joshua Kistler, Sophia Rehm, Katharina Rentsch, Stefan Borgwardt, Christian Cajochen, Carolin Reichert

**Affiliations:** Centre for Chronobiology, Psychiatric Hospital of the University of Basel, Basel, Switzerland; Transfaculty Research Platform Molecular and Cognitive Neurosciences, University of Basel, Basel, Switzerland; Neuropsychiatry and Brain Imaging, Psychiatric Hospital of the University of Basel, Basel, Switzerland; Institute of Pharmacology and Toxicology, University of Zürich, Zürich, Switzerland; Sleep & Health Zürich, University Center of Competence, University of Zürich, Zürich, Switzerland; F. Hoffmann-La Roche Ltd, Basel, Switzerland; Laboratory Medicine, University Hospital Basel, Basel, Switzerland

## Abstract

To enhance wakefulness, daily consumption of caffeine in the morning and afternoon is highly common. However, it is unknown whether such a regular intake pattern affects timing and quality of wakefulness, as regulated by an interplay of circadian and sleep-homeostatic mechanisms. Thus, we investigated the effects of daily caffeine intake and its withdrawal on circadian rhythms and wake-promotion in 20 male young habitual caffeine consumers. We applied a double-blind, within-subject design with a caffeine (150 mg, 3 x daily), a placebo, and a withdrawal condition each lasting ten days. Starting on day nine of treatment, salivary melatonin and cortisol, evening nap sleep, as well as sleepiness and vigilance performance throughout day and night were quantified during 43 h under controlled laboratory conditions. Neither the time course of melatonin (i.e., onset, amplitude, or area under the curve) nor the time course of cortisol were significantly affected by caffeine intake or its withdrawal. During withdrawal, however, volunteers reported increased sleepiness, showed more attentional lapses, as well as polysomnography-derived markers of elevated sleep propensity in the late evening compared to both, placebo and caffeine conditions. Thus, the typical timing of habitual caffeine intake in humans may not necessarily shift circadian phase nor lead to clear-cut benefits in alertness. The time-of-day independent effects of caffeine withdrawal suggest an adaptation to the substance, presumably in the homeostatic aspect of sleep-wake regulation.

## Introduction

Caffeine is the most commonly consumed psychoactive substance in the world [1]. Around 80% of the worldwide population consume caffeine regularly on a daily basis [2] and intake is increasing in terms of daily dosages and earlier age of regular substance intake [3]. Caffeine containing aliments, e.g. coffee, tea, soda drinks, and chocolate [1], are used since centuries to modulate sleep and wakefulness [4].

Timing, quality, and quantity of sleep and wakefulness are regulated by the interplay of a homeostatic and circadian process [5]. Caffeine interferes with sleep homeostasis by antagonizing adenosine receptors [6], a proposed mediator of the increase of homeostatic sleep pressure during time spent awake and its decrease during sleep [7]. By blocking the A_1_ and A_2A_ adenosine receptors [1], which are expressed in wide-spread areas of the human central nervous system [8,9], acute caffeine administration reduces the effects of sleep pressure, as mirrored in reduced sleepiness [10], improved behavioral performance [11], and dampened sleep depth during nighttime sleep [6], particularly when sleep pressure is high [3,12].

Furthermore, evidence accumulates that caffeine also impacts on circadian rhythms, in humans usually assessed by changes in salivary melatonin levels. Acute caffeine intake in the evening and at night has been shown to delay the onset of melatonin secretion [13] and decrease nighttime melatonin levels [14,15]. However, evening intake of caffeine is not common in the society nowadays [16]. Considering the average half-life of caffeine with a duration of around 4 h [12], the question arises whether caffeine-induced circadian effects also occur when consumption is timed to morning and/or afternoon, as observed in habitual caffeine consumers [16].

Besides the timing of caffeine intake, the duration of prior repeated daily use moderates the impact of caffeine-induced changes on sleep-wake regulation. There is evidence that consumers develop tolerance to the substance already after several days, such that effects of a particular dose of caffeine, for instance on sleep [17] or alertness [10] become weakened. However, in line with a recent study in animals [18], it has also been shown that continuous hourly caffeine intake over four weeks strengthens circadian wake-promotion, as indicated by a reduced ability to sleep prior to habitual bedtime [19]. Interestingly, timing of melatonin secretion was not shifted by the long-term treatment. An open question remains, whether the absence of phase shifts can be traced back to the continuous timing of caffeine administration around the entire circadian cycle or due to neuroadaptations in response to long-term treatment.

One of the typical indicators of neuronal and systemic alterations in response to long-term caffeine use is the occurrence of withdrawal symptoms when intake is ceased [20]. Caffeine withdrawal symptoms include decreased alertness [10,21], impaired cognitive performance [10,21], and changes in waking electroencephalogram (EEG) such as enhanced theta power [22], starting 12 to 24 h after last caffeine intake with peak intensity between 20 to 51 h and a maximal duration of nine days [20]. Based on changes in adenosine-signaling [23], caffeine withdrawal thus constitutes states of low arousal which may help triggering the maintenance of daily caffeine intake in habitual consumers [20]. However, to our best knowledge the impact of caffeine-withdrawal on human circadian sleep-wake regulation has not yet been examined.

Thus, we investigated the effects of daily daytime caffeine consumption and its withdrawal on human waking performance, circadian rhythms, and wake-promotion. To establish both tolerance and withdrawal, and to enable a comparison to a withdrawal-free baseline, caffeine and placebo, respectively, were administered over ten days in a crossover design with three conditions (caffeine, placebo, and withdrawal). Starting at day nine of treatment, sleepiness, and vigilance performance was assessed as well as salivary melatonin, cortisol, and nap sleep during high circadian wake-promotion within a 43h-laboratory protocol under controlled light, posture, and meal intake.

## Materials and Methods

The present study was approved by the Ethics Committee northwest/central Switzerland (EKNZ) and conducted in accordance with the declaration of Helsinki. All volunteers provided written informed consent and received a monetary compensation for study participation.

### Volunteers

In total, 179 healthy, habitual caffeine consumers underwent a thorough screening procedure. Exclusion criteria comprised age < 18 or > 35 years, body mass index (BMI) < 18 or > 26, drug dependency, shiftwork within 3 months prior to study admission, transmeridian travels within one month prior to study, extreme chronotype (Morningness-Eveningness Questionnaire [24], score < 30 and > 70) and poor sleep quality (Pittsburgh Sleep Quality Index [25], score > 5). Volunteers were included when habitual daily caffeine intake was between 300 and 600 mg, assessed with a survey tool [26] adapted to the caffeine content according to [12]. Twenty-nine volunteers were invited for a habituation night and a physical examination by a physician in charge to exclude poor sleep efficiency (SE < 70%), clinical sleep disturbances (apnea index > 10, periodic leg movements > 15/h), and chronic or debilitating medical conditions. Demographic characteristics of the 20 participants who completed all three conditions can be found in the supplementary materials (Table S1).

### Design and Protocol

The protocol is illustrated in Figure 1A. A double-blind, crossover study comprising three conditions was conducted: a caffeine, a placebo, and a withdrawal condition. This within-subject design was chosen to reduce variance in the data due to expectancies or inter-individual variability in caffeine metabolism. For calculation of sample size and pseudo-random allocation of volunteers to the order of the three conditions, see supplementary materials (Table S2). In each condition, participants ingested gelatin capsules over ten days three times daily (+45 min, +255 min, +475 min after wake-up), containing either placebo (mannitol, Hänseler AG, Herisau, Switzerland) or caffeine (150 mg, Hänseler AG, Herisau, Switzerland). Participants were instructed to refrain from caffeinated beverages and food. Compliance was verified by assessing caffeine metabolites from fingertip sweat collected prior to habitual bedtime (see supplementary materials). The length of the treatment of ten days was based on the maximum duration of withdrawal symptoms [20], occurring during placebo treatment in habitual consumers. Timing and dose were based on an earlier study investigating tolerance to caffeine and its withdrawal [10].

**Figure 1.**
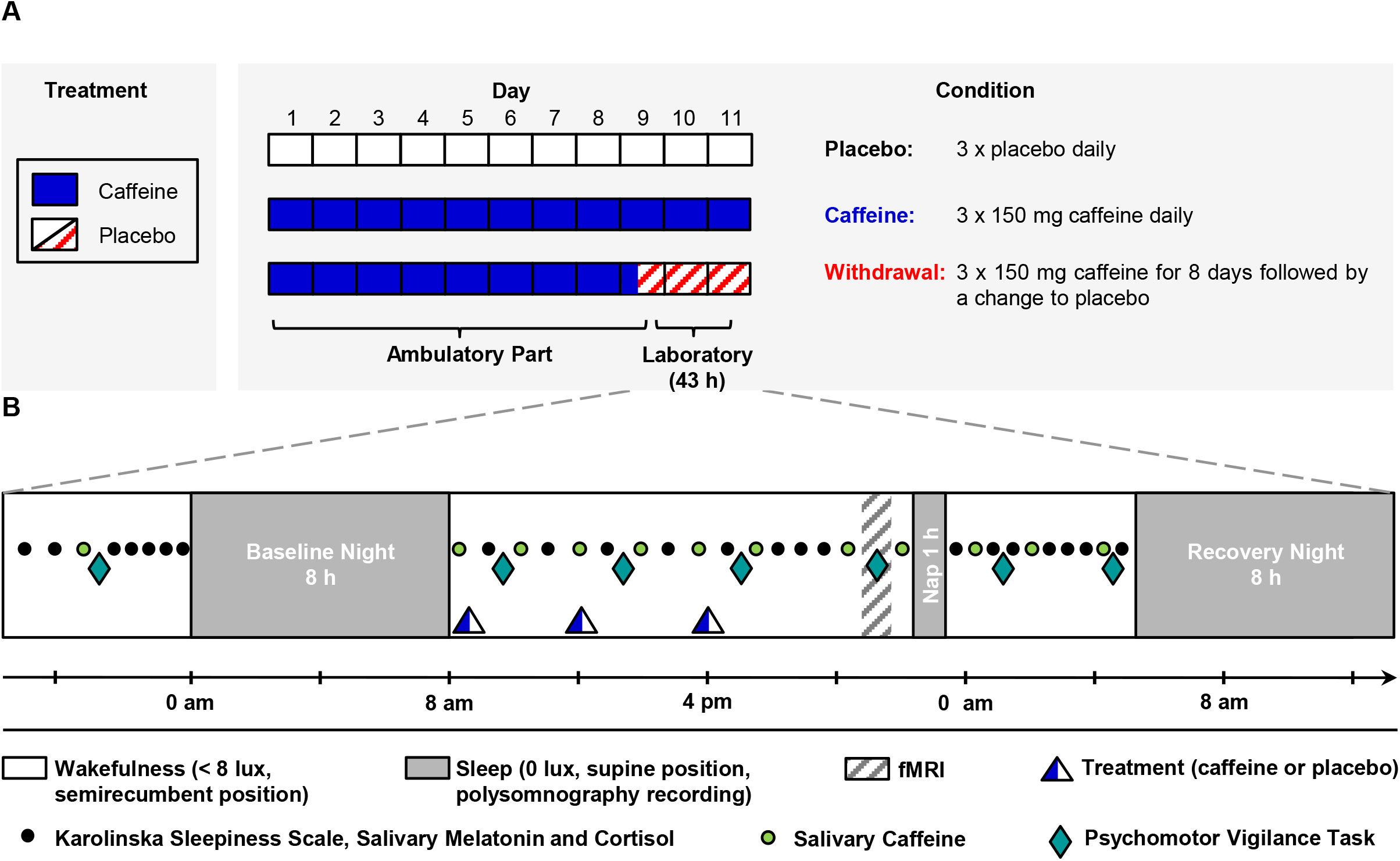
Schematic illustration of study protocol. **A** Each volunteer participated in a placebo, a caffeine, and a withdrawal condition, each comprising a 43-h laboratory stay, preceded by an ambulatory part. In each condition either caffeine or placebo capsules were administered three times daily. **B** The laboratory stay started in the evening of day nine with assessments of subjective sleepiness, vigilance performance, salivary melatonin, cortisol, and caffeine levels, which were continued on the next day after an 8-h baseline night. In the evening of day ten, 14.5 h after wakeup, a 1-h nap was scheduled during a time of high circadian wake-promotion [33,34] followed by 5.75 h of scheduled wakefulness and an 8-h recovery night.

The caffeine and placebo condition included one type of pill exclusively (caffeine or placebo, respectively). The withdrawal condition comprised a switch from caffeine to placebo pills in the late morning of day nine (+255 min after wake-up). In scheduling the caffeine to placebo switch to the late morning, a coincidence of the peak of withdrawal symptoms (after around 35 h after last caffeine intake) with a window of high circadian wake-promotion was aimed for. Dim light melatonin onset (DLMOnset) was used as marker of circadian timing.

Each condition started with an ambulatory part of nine days during which participants ingested capsules according to the regimen described above. In addition, participants kept a fixed sleep-wake rhythm (within +/− 30 min of self-selected bedtime, time in bed 8 h, no naps) verified by wrist actimetry (Actiwatch, Cambridge Neurotechnology, Cambridge, United Kingdom). Compliance to drug abstinence was checked prior to each laboratory part by a urinary toxicology screen (AccuBioTech Co. Ltd, Beijing, China).

In the evening on day nine of treatment, 5.5 h before individual habitual bedtime, a 43-h laboratory part started, schematically illustrated in Figure 1B. Volunteers were accommodated in single apartments, isolated from external time cues and communication was restricted to team members. Additionally, light conditions (< 8 lux), posture, and meal intake was controlled.

### Salivary caffeine

Saliva samples to quantify caffeine levels were collected in intervals of approximately 2 h during scheduled wakefulness during the laboratory stay. Subsequently, caffeine levels were analyzed with liquid chromatography coupled to tandem mass spectrometry. One dataset was not included due to its non-availability.

### Neurobehavioral assessments

Subjective sleepiness was assessed regularly with the Karolinska Sleepiness Scale (KSS) [27] every 30 to 60 min during scheduled wakefulness. For analyses, values were binned to 4-h intervals. Vigilance performance was measured by a visual 10-minute psychomotor vigilance task (PVT) [28], every 4 h during scheduled wakefulness. Volunteers were instructed to focus on a white cross displayed on a black screen and to respond as fast as possible by a key press as soon as a millisecond counter appeared. The inter-stimulus interval was randomized between 2 and 10 s. Here, we focus on the number of lapses (reaction time > 500 ms), the most sensitive measure for the interaction of caffeine with both sleep pressure and circadian phase [19]. As one of the tests was conducted in a magnetic resonance scanner with a compatible button box, lapses were z-transformed before analyses according to this change in environment.

### Salivary melatonin and cortisol

Saliva samples were collected regularly in intervals of 30 to 60 min. For handling, see supplementary materials. Melatonin and cortisol levels were detected using a direct double-antibody radio immunoassay [29] and an enzyme-linked immunosorbent assay (ALPCO, Salem, NH, USA), respectively. For analyses, data were collapsed into bins of 1.5 h.

Four datasets were excluded from melatonin analyses due to insufficient data quality (placebo condition: one; caffeine condition: two; withdrawal condition: one). For analyses of melatonin, data were resampled every minute by applying linear interpolation. Subsequently, a bimodal skewed baseline cosine function (BSBCF) curve [30] was fitted to the data based on [31] with the modified cost function proposed in [32]. Goodness of fit (R^2^) was acceptable for all data sets (indicator > 0.6) except for two volunteers (placebo condition: one; caffeine condition: one). DLMOnset and dim light melatonin offset (DLMOffset) were determined for the fitted BSBCF curve applying a threshold of 0.1 of its amplitude which was defined by the difference between peak to baseline levels [30]. In order to estimate condition-specific changes in the melatonin profile, the amplitude and the area under the curve (AUC) were calculated including samples following wake-up at day ten of treatment.

### Polysomnography during nap sleep

To test caffeine-induced differences in circadian wake-promotion, EEG was recorded during a one hour nap episode in the evening, starting 14.5 h after wake-up. It has repeatedly been shown that the ability to sleep is lowest in the evening [33,34], mirroring maximal wake-promoting strength at the end of the day. Recorded EEG data were visually scored according to [35], blind to the condition, yet slow-wave sleep (SWS) was further classified into stages 3 and 4 [36]. For details on recording procedure and scoring, see supplementary materials.

Total sleep time (TST) was calculated as sum of sleep stages 1, 2, 3, 4, and rapid eye movement (REM) sleep. Sleep efficiency (SE) was calculated as TST divided by time in bed. SWS comprises the sum of sleep stage 3 and 4. Sleep latency 1 and 2 were defined as latency to the first occurrence of sleep stage 1 and 2, respectively. Non-REM (NREM) sleep was calculated as sum of stage 2, 3 and 4. Duration of REM sleep was not analyzed as most participants (N = 16, 93%) did not reach this sleep state. To test condition-specific differences in the time course of nap sleep, TST and SWS was collapsed into 5-min time bins.

Spectral analysis of NREM sleep was conducted using a Fast Fourier Transformation (FFT) on 4-s time windows (hamming, 0% overlapped) resulting in 0.25 Hz bins. Frequency bins from 0.5 – 32 Hz of NREM sleep, recorded from frontal derivations (F3, F4), were analyzed. Data were log-transformed and collapsed into 1 Hz bins. Note that condition-specific analyses are based on a reduced number of datasets because ten participants did not initiate NREM sleep in at least one of the three conditions (four participants in placebo, six participants in caffeine, none in the withdrawal conditions).

### Statistical Analyses

Data analyses were conducted with the statistical analyses software (SAS Institute, Cary, NC, USA) version 9.4 using mixed model analysis of variance (PROC MIXED) with *subject* as a random factor and the two repeated factors *condition* (three levels: placebo, caffeine, and withdrawal) and *time* (levels differ per variable). To account for correlations between adjacent points within *time*, we used AR(1) as covariance structure (i.e., autoregressive (1)). In analyses without the factor *time*, we assumed CS (i.e., compound symmetry) to model the covariance structure most properly. Degrees of freedom were adjusted based on Kenward-Roger [37]. *P*-levels of post-hoc comparisons, derived from the LSMEANS statement, were corrected for multiple comparisons with the Tukey-Kramer method. Data of one participant in the caffeine condition have been excluded from all analyses due to incompliance of the treatment.

## Results

### Caffeine levels

As expected, caffeine levels were higher in the caffeine condition compared to both the withdrawal and placebo condition (main effect of condition: *F*_2,121_ = 185.16; *p* < 0.0001; post-hoc tests: *p* < 0.0001) while levels during withdrawal were increased compared to the placebo condition (post-hoc tests: *p* = 0.035; mean ± SD: placebo: 18.52 ± 84.80 ng/ml; caffeine: 3024.18 ± 2163.68 ng/ml; withdrawal: 296.81 ± 683.42 ng/ml). As illustrated in Figure S3, this general pattern was modulated by time (interaction condition × time (*F*_22,237_ = 2.76; *p* < 0.0001), mirroring both a decrease of caffeine levels during withdrawal condition and an increase in the caffeine condition after administration of treatment.

### Subjective Sleepiness and Vigilance Performance

The time course of subjective sleepiness values for each of the three conditions is illustrated in Figure 2A and the number of lapses on the PVT in Figure 2B.

**Figure 2.**
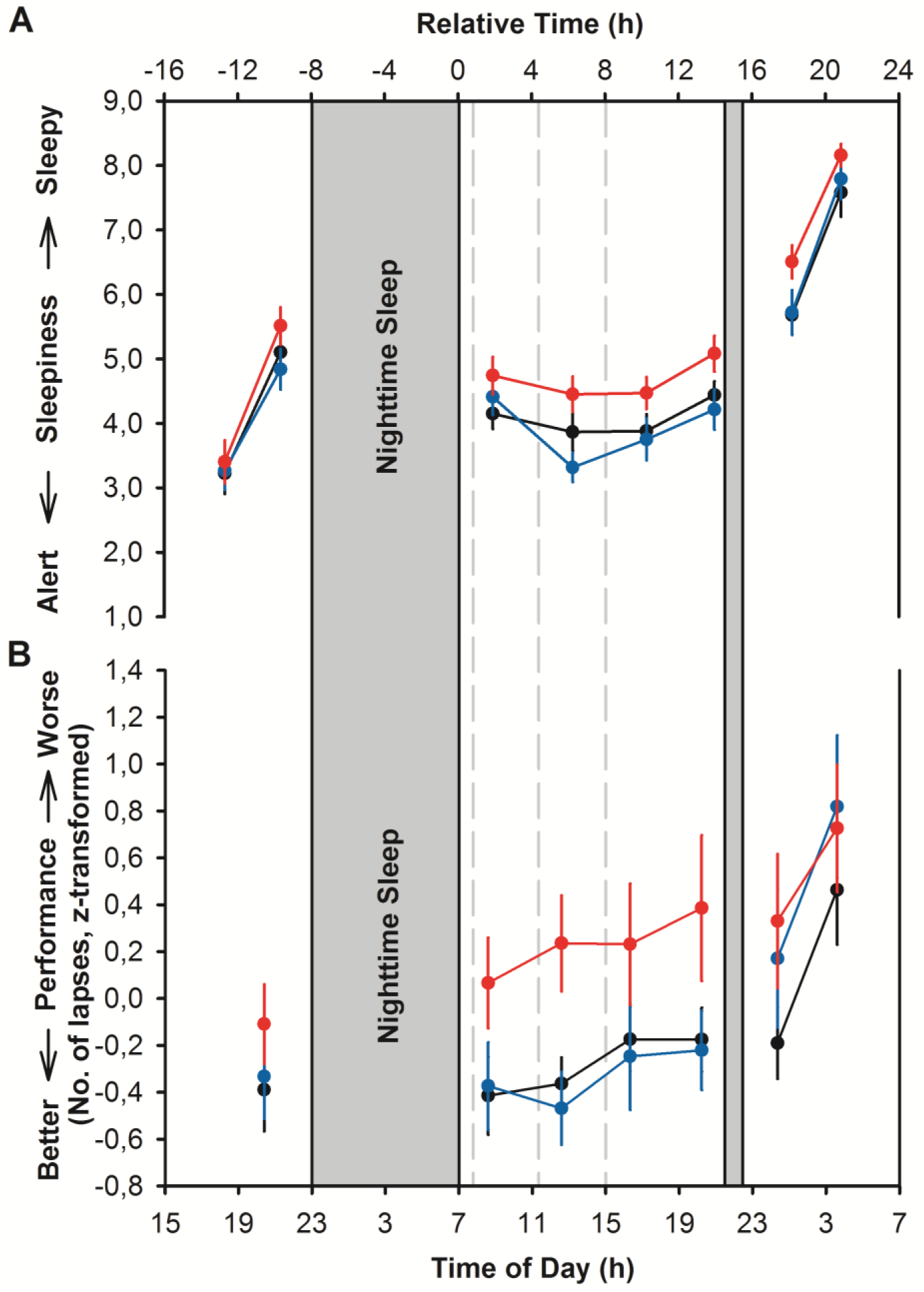
Time course of subjective sleepiness (A) and attentional lapses assessed during PVT (B) in the placebo (black), caffeine (blue), and withdrawal (red) condition across 43-h (means ± standard errors). Pill administrations (caffeine or placebo) are depicted with dashed lines. Both subjective sleepiness and vigilance performance were impaired after the abrupt cessation of caffeine compared to placebo and daily caffeine intake.

The results of the mixed model analysis of variance indicated higher subjective sleepiness in the withdrawal condition compared to the placebo and caffeine conditions (main effect of condition: *F*_2,58_ = 9.71; *p* < 0.001, post-hoc tests: *p* < 0.01; mean ± SD: placebo: 4.72 ± 1.78; caffeine: 4.66 ± 1.91; withdrawal: 5.27 ± 1.81). A significant main effect of time (*F*_7,127_ = 54.62; *p* < 0.0001) confirmed a diurnal profile with higher sleepiness during the biological night compared to daytime.

The analyses of the number of lapses on the PVT yielded a significant main effect of condition (*F*_2,51_ = 6.66; *p* = 0.0027; post-hoc tests: *p* < 0.01) revealing more lapses during the withdrawal condition compared to the placebo and caffeine conditions (mean ± SD: placebo: −0.18 ± 0.77; caffeine: −0.09 ± 1.04; withdrawal: 0.27 ± 1.11). A significant main effect of time (*F*_6,98_ = 7.55; *p* < 0.0001) indicated a typical profile of more lapses during the night compared to daytime.

### Melatonin and Cortisol

In the analyses of melatonin levels, only the main effect of the factor time (*F*_17,290_ = 33.44; *p* < 0.0001) was significant, confirming a diurnal profile of higher melatonin levels during the night compared to morning (Figure 3A). Neither DLMOnset (*F*_2,33_ = 1.16; *p* = 0.325) nor DLMOffset (*F*_2,35_ = 1.32; *p* = 0.280) significantly differed between conditions. Similarly, the effect of condition on amplitude (*F*_2,34_ = 0.57; *p* = 0.573) and AUC (*F*_2,34_ = 1.77; *p* = 0.185) did not reach significance. For means and standard errors of the melatonin outcomes, see Table S3.

**Figure 3.**
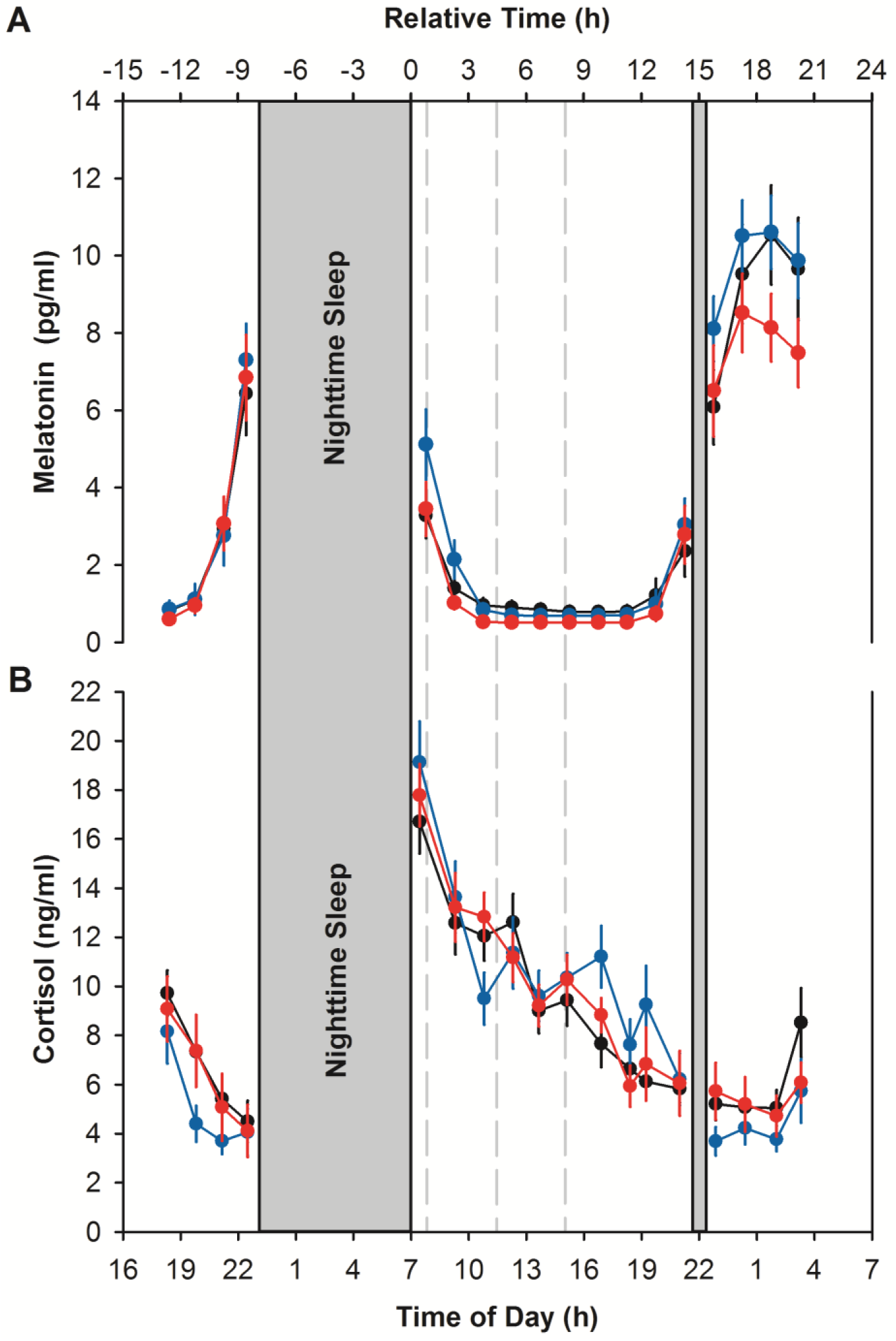
Time course of salivary melatonin (A) and cortisol (B) across the 43-h laboratory stay. The means ± standard errors are depicted for placebo (black), caffeine (blue), and withdrawal conditions (red) and pill administrations (caffeine or placebo) are indicated with dashed lines. The melatonin profile depicts the fitted data. The typical course of both melatonin and cortisol across a day is clearly visible, however being independent of the treatment.

Analyses of cortisol levels did not show significant differences between conditions (*F*_*2,129*_ = 0.43; *p* = 0.653). However, a significant main effect of the factor time (*F*_17,303_ = 31.66; *p* < 0.0001) demonstrated a normal diurnal pattern with higher cortisol levels in the morning and lower levels during the night. The time course of salivary cortisol levels is illustrated in Figure 3B.

### Evening Nap Sleep

A summary of sleep variables per condition and results of the statistical analyses are presented in Table 1. TST and SWS were longer, and SE was higher during withdrawal compared to placebo and caffeine conditions (post-hoc tests: *p* < 0.05). Furthermore, sleep latency to sleep stage 1 and stage 2 were shorter during withdrawal compared to placebo and caffeine conditions (post-hoc tests: *p* < 0.05). Generally, initiation of NREM sleep was less frequent in the caffeine and placebo condition compared to withdrawal (no NREM sleep in placebo: N = 4, caffeine: N = 6, and withdrawal condition: N = 0, Cochrans Q-test: *p* < 0.05).

**Table 1.** Sleep parameters derived from visual scoring assessed during evening nap sleep.

| Sleep Parameter | Placebo | Caffeine | Withdrawal | Factor Condition |
| --- | --- | --- | --- | --- |
| TST (min) | 31.63 ± 3.90 | 24.26 ± 4.55 | 43.88 ± 2.83<sup>a</sup> | $F(2,37) = 9.64, p < 0.001$ |
| SE (%) | 52.71 ± 6.50 | 40.44 ± 7.58 | 73.12 ± 4.72<sup>a</sup> | $F(2,37) = 9.64, p < 0.001$ |
| Stage 1 (min) | 4.95 ± 0.73 | 4.66 ± 0.83 | 5.80 ± 0.85 | $F(2,38) = 0.67, p = 0.519$ |
| Stage 2 (min) | 12.10 ± 1.97 | 9.82 ± 2.22 | 15.30 ± 1.57<sup>b</sup> | $F(2,37) = 3.85, p = 0.030$ |
| Stage 3 (min) | 6.53 ± 1.48 | 5.74 ± 1.73 | 10.20 ± 1.96<sup>b</sup> | $F(2,37) = 4.09, p = 0.025$ |
| Stage 4 (min) | 6.80 ± 2.16 | 3.74 ± 2.01 | 11.48 ± 2.69<sup>b</sup> | $F(2,37) = 6.06, p = 0.005$ |
| SWS (min) | 13.33 ± 2.82 | 9.47 ± 2.70 | 21.68 ± 3.00<sup>a</sup> | $F(2,37) = 8.22, p = 0.001$ |
| NREM (min) | 25.43 ± 4.01 | 19.29 ± 4.27 | 36.98 ± 3.34<sup>a</sup> | $F(2,37) = 8.87, p < 0.001$ |
| SL1 (min) | 25.70 ± 3.94 | 28.51 ± 4.57 | 12.09 ± 2.11<sup>a</sup> | $F(2,37) = 7.59, p = 0.002$ |
| SL2 (min) | 31.16 ± 4.11 | 37.94 ± 4.51 | 18.39 ± 2.87<sup>a</sup> | $F(2,37) = 9.80, p < 0.001$ |

Values are for means ± standard errors. TST = Total sleep time, SE = Sleep efficiency, SWS
= Slow-wave sleep (stage 3 + 4), NREM = Non-rapid eye movement sleep, SL1 = Sleep
latency to sleep stage 1. SL2 = Sleep latency to sleep stage 2.
<sup>a</sup> $p < 0.05$ compared to placebo and caffeine condition
<sup>b</sup> $p < 0.05$ compared to caffeine condition

In a next step we were interested whether TST and SWS accumulate differently depending on treatment. As depicted in Figure 4, particularly in the beginning of the nap, TST and SWS accumulated faster during the withdrawal condition compared to caffeine and placebo conditions (interaction condition × time for TST: *F*_*22,354*_ = 5.40; *p* < 0.0001; and SWS: *F*_*22,354*_ = 3.82; *p* < 0.0001).

**Figure 4.**
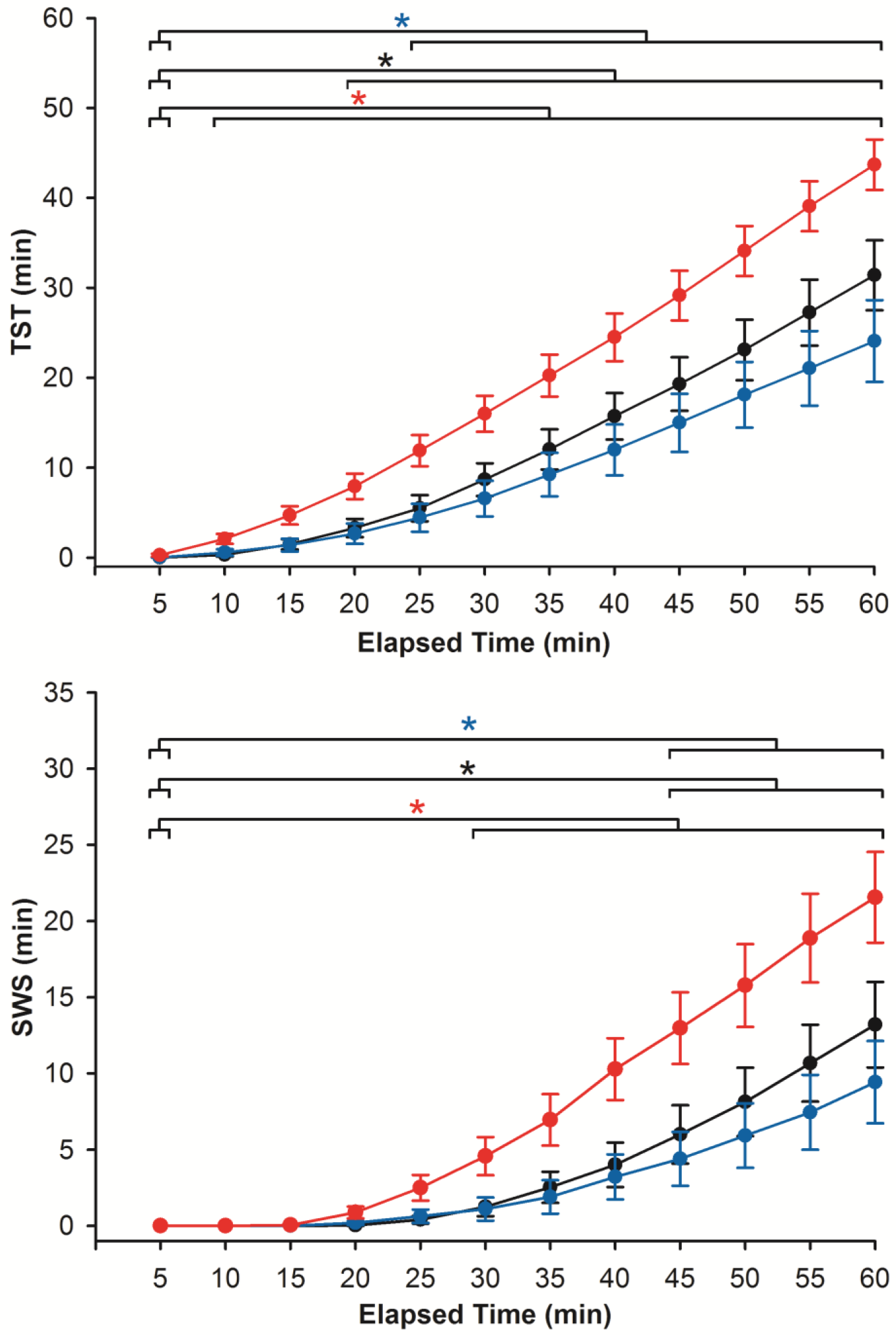
Accumulation curves of TST and SWS during evening nap sleep. Data were collapsed into bins of 5 min and accumulated across the one hour. Means and standard errors are represented for placebo (black), caffeine (blue), and withdrawal (red). Asterisks mark significant differences to the first bin within the same condition (*p*_*all*_ < 0.05).

In a final step, differences in spectral power density between conditions were analyzed. There were no significant differences in spectral power density between caffeine and placebo. However, spectral power density was reduced in the sigma range during the withdrawal compared to placebo condition (15 Hz: *F*_*2,27*_ = 3.48; *p* = 0.045: 16 Hz: *F*_*2,28*_ = 3.15*; p* = 0.059).

## Discussion

The aim of the present study was to examine the effects of daily caffeine intake and its cessation on circadian timing, wake-promotion, and the course of neurobehavioral indices in habitual consumers under entrained conditions. Making use of a carefully controlled within-subjects design, we focused on the effects of repeated daily consumption in the morning and afternoon, as this is the typical pattern in the society nowadays. Under these conditions, neither caffeine consumption nor its cessation affected the diurnal profile of melatonin or cortisol secretion. However, for more than 20 h, cessation of intake induced a state of withdrawal, characterized by higher subjective sleepiness, impaired vigilance performance, and a higher drive to sleep even at a phase of high circadian wake-promotion. It is concluded that daily daytime intake of caffeine does not strongly affect human circadian rhythms under entrained conditions. Still, it induces adaptations, potentially at the level of the adenosine receptors, which come to light as a state of reduced arousal and alertness as soon as caffeine intake is ceased.

In animal studies, long-term treatment with caffeine leads to a different timing of circadian rest-activity rhythms, specifically to a lengthening of circadian period under constant lighting conditions [38,39]. Similar observations in unmasked human rest-activity cycles are missing so far. However, under conditions of forced desynchronization between the timing of circadian rhythms and sleep-wake cycles, circadian plasma melatonin rhythms were not altered in response to hourly administration of caffeine over four weeks [19]. The results of the present study add that – under entrained conditions – the timing of both melatonin and cortisol does not necessarily change by daily caffeine consumption in the morning, midday, and/or afternoon, a temporal pattern which is observed in around 80% of consumers [16]. Apart from statistics on the group’s average, visual inspection however revealed a delay in DLMOffset during caffeine in more than half of the volunteers (N = 13, see supplementary materials). Thus, interindividual differences in response to the stimulant [40] as well as the small sample size might have hampered statistical condition-specific differences at *p* < 0.05. Importantly, based on the absence of a clear-cut shift in circadian timing in the withdrawal condition, it seems unlikely that the participants might have developed tolerance to the phase-shifting effects of the drug. Moreover, recent animal studies on repeated caffeine treatment indicate no [38,41] or only slight [18] caffeine-induced phase delays in locomotor activity under normal light-dark cycles. Taken together the available evidence indicates that the effects of repeated daytime caffeine intake on human circadian timing under entrained conditions seem to be small or absent.

Most likely, however, the effects of caffeine on circadian rhythms depend on the time of intake. So far, a delay [13] or reduction in melatonin [14] in humans, either abstinent [13] or potentially under withdrawal [14], was specifically induced after a caffeine treatment in the evening or at night. In contrast, repeated caffeine intake in the morning did not successfully entrain three blind patients [42]. Furthermore, a recent animal study under constant conditions suggests that caffeine treatment does only potentiate light-induced phase shifts when given at the end of the active phase or during rest, but not at start of the active phase [43]. Together with the present results, the evidence suggests that the circadian system seems to be particularly sensitive to caffeine when given at the end of the biological day. Future studies might disentangle circadian and sleep-homeostatic contributions to this effect. Independent of time-of-day, we observed clear-cut effects induced by caffeine withdrawal. In line with earlier studies [20], the acute challenge of cessation from caffeine was associated with signs of increased sleep pressure, such as increased subjective sleepiness and worse vigilance performance during day and nighttime as well as a faster initiation of NREM sleep even at a time of high circadian wake-promotion. Thus, the preceding repeated presence of caffeine might have induced compensatory adaptations at the neuronal level [23], which modulate the stimulatory effects of caffeine and underlie the effects of withdrawal as soon as consumption is stopped. Several changes have been associated with long-term caffeine intake, e.g. upregulation of adenosine receptors [44–47], increased plasma adenosine concentrations [48] or modulations in the function of adenosine heteromers [49]. These neuronal changes in the adenosinergic system might alter the homeostatic sleep need. In the present study, an increased sleep pressure experienced during caffeine withdrawal might have overruled the circadian drive for wake-promotion in the evening, a phenomenon which has already been shown after sleep restriction in humans [50].

In contrast to the clear-cut symptoms of caffeine withdrawal in behavior, we do not have any indication for a significant difference in either of the measured variables during daily caffeine consumption compared to placebo. Importantly our study was designed to focus on the effects of caffeine after a certain period of repeated intake under normal sleep-wake conditions. In earlier studies that showed a caffeine-induced sleep disruption during circadian wake-promotion after continuous daily intake, measurements were taken under relatively high levels of sleep pressure (i.e., after 25 h [51] or 28 h [19] of wakefulness). Interestingly, reviews indeed suggest that the stimulating properties of caffeine are most prominently under high sleep pressure such as after sleep deprivation or sleep restriction [3,12]. Therefore, we may not exclude that caffeine intake would have induced sleep disruption and alertness under a longer duration of wakefulness as was applied in the present study. However, studies controlling for withdrawal reversal, similarly applied in our study, failed to show a caffeine-induced improvement in performance in sleep-restricted subjects [52,53], indicating that improvements by caffeine cannot solely be explained by sleep-wake-history but probably also depend on preceding caffeine intake. Applying repeated caffeine administrations and a wash-out period of nine days prior to each assessment phase, potential effects in the caffeine condition deriving from withdrawal reversal and carry-over effects can likely be excluded.

Moreover, one might argue that the lack of improvement in subjective alertness and performance during caffeine intake compared to placebo condition is due to a floor effect. In other words, the low sleepiness and high performance level occurring during the placebo condition did not leave much room for additional improvement by caffeine. However, our measurements took place also during the biological night, in which we observed the typical nighttime decrease in alertness and performance. As we did not observe a significant caffeine-induced improvement under these conditions, our results suggest that the effects of daytime caffeine intake on alertness and performance are either short-lasting, small or not present under conditions of habitual daily caffeine intake.

Finally, it has been suggested that the dose-response of caffeine on performance asymptotes at around 200 mg of caffeine [54]. However, the effects are ambiguous with studies showing performance improvements with higher caffeine doses [54,55]. Furthermore, in habitual consumers, as included for the present study, doses of 400 mg have been shown to be required to induce certain performance benefits [56]. Nevertheless, habitual consumption level might be an indicator for individuals who are less sensitive to caffeine [57] Thus, the question arises whether the results indicate no-effects as a consequence of the self-selected study sample of habitual consumers or mirror tolerance development. As the present study design does not include a condition assessing the effects of acute caffeine intake after long-term abstinence, we cannot provide a strict measure of tolerance by comparing acute effects of caffeine with the effects after daily treatment in the same individuals. However, there are convincing indicators for tolerance to occur roughly after 3-5 days of habitual caffeine intake [58] with the potential of complete [59] or partial tolerance [60]. Moreover, the effects yielded when caffeine was ceased provide evidence for an adaptation to the daily exposure of caffeine in waking performance, however circadian timing and amplitude remains mainly unaffected.

Taken together, this is the first study investigating the impact of habitual caffeine consumption on human circadian rhythms under entrained conditions. The study was designed to focus on the effects of a typical pattern of caffeine consumption with daily intake in the morning and afternoon. We provide first evidence that this type of exposure to the stimulant does not considerably shift circadian markers such as melatonin and cortisol nor does it lead to an increased wake-promotion in the evening. However, the acute challenge of cessation from caffeine was associated with signs of increased sleep pressure. Together, our data point to an adaptation of waking-performance to habitual exposure to the stimulant while circadian markers remain fairly stable. These mechanisms of both adaptation and robustness might enable normal sleep-wake states during constant supply of a stimulating agent in the central nervous system.

## Acknowledgements

We thank our interns Andrea Schumacher, Laura Tincknell, Sven Leach and all the study helpers for their great help in data collection, Claudia Renz for the analyses of melatonin and cortisol, Dr. Martin Meyer and Dr. Helen Slawik for the physical examinations, and all our volunteers for study participation.

