## Supplemental Material for "Caffeine-dependent changes of sleep-wake regulation: evidence for adaptation after repeated intake"

Description of study sample

**Table S1.** Demographic characteristics of study sample.

| Demographics | Mean ± SD |
|---|---|
| Age (years) | 26.4 ± 4.0 |
| Habitual daily caffeine intake (mg) | 478.1 ± 102.8 |
| Body Mass Index (kg/m$^2$) | 22.7 ± 1.4 |
| Morningness-Eveningness Questionnaire | 52.8 ± 8.7 |
| Munich ChronoType Questionnaire | 4.2 ± 0.7 |
| Epworth Sleepiness Scale | 3.6 ± 3.4 |
| Pittsburgh Sleep Quality Index | 2.8 ± 1.4 |
| Beck Depression Inventary-II | 1.4 ± 2.3 |

Rationale for sample size

Calculation of sample size was done with G*Power 3.1 [1], planning the calculation of an ANOVA for repeated measures (three conditions) with an accepted α error probability = 0.05 and power (1-β) = 0.8. For an estimation regarding circadian phase, the assumed correlation between measures ($r$ = 0.7) was based on a previous study, in which melatonin was assessed in two differential sleep pressure conditions in young healthy volunteers after seven days of a fixed sleep-wake cycle [2]. The expected effect size ($f^2$ = 0.26) is assumed on the basis of [3] investigating the influence of caffeine on melatonin levels.

Frequency of order of conditions

Volunteers were pseudo-randomly allocated to the order of the three conditions based on random permutation controlling for block size.

**Table S2.** Number of participants per order of conditions.

| Caffeine - Placebo - Withdrawal | Caffeine - Withdrawal - Placebo | Withdrawal - Placebo - Caffeine | Withdrawal - Caffeine - Placebo | Placebo - Caffeine - Withdrawal | Placebo - Withdrawal - Caffeine |
|---|---|---|---|---|---|
| N = 3 | N = 3 | N = 4 | N = 3 | N = 4 | N = 3 |

Caffeine levels during ambulatory phase

In order to verify volunteers' compliance to the regimen prior to laboratory admission, samples containing fingertip sweat were collected approximately 8 h after the last pill intake on days one to eight and 5 h after the last pill intake on day nine of treatment. Subsequently, caffeine levels were analyzed by triple-quadrupole mass spectrometry.

A significant main effect of condition ($F_{2,52}$ = 21.70; $p$ < 0.0001) confirms adherence to the protocol by increased caffeine levels during the caffeine and withdrawal conditions compared to the placebo condition ($p_{all}$ < 0.0001), depicted in Figure S1.

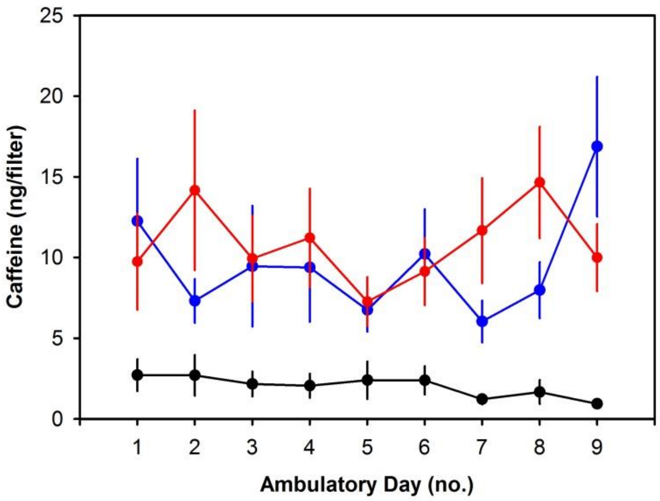


**Figure S1.** Time course of caffeine levels collected during the ambulatory phase in the placebo (black), caffeine (blue), and withdrawal (red) condition confirming volunteers' compliance to the regimen.

Handling and analyses of saliva samples

Saliva samples were collected regularly in intervals of 30 to 60 min under dim light conditions (< 8 lux). Following collection, samples were stored at maximum -5°C, later centrifuged (3000 rpm for a duration of 10 min), and subsequently stored at -24°C until data analyses. Melatonin and cortisol levels were analyzed by a trained staff member (at Bühlmann Laboratories AG, Schönenbuch, Switzerland) using a direct double-antibody radio immunoassay [4] and an enzyme-linked immunosorbent assay (ALPCO, Salem, NH, USA), respectively.

To test caffeine-dependent shifts in circadian timing, we analyzed condition-specific differences in dim light melatonin onset (DLMOnset) and dim light melatonin offset (DLMOffset). In order to account for slightly different bedtimes between conditions in three volunteers (twice: +30 min in caffeine compared to placebo and withdrawal condition; once: -30 min in placebo compared to caffeine and withdrawal condition), we further tested condition-specific effects in phase angle (difference of bedtime – DLMOnset). However, phase angle did not significantly differ among the three conditions on day nine or on day ten of treatment ($p_{all} > 0.2$). A summary of melatonin parameters per condition and results of the statistical analyses is depicted in Table S3.

**Table S3.** Melatonin parameters assessed on day ten of treatment.

| Melatonin Parameter | Placebo | Caffeine | Withdrawal | Factor Condition |
|---|---|---|---|---|
| DLMOnset (h) | 21.74 ± 0.33 | 21.15 ± 0.18 | 21.60 ± 0.31 | $F(2,33) = 1.16$, $p = 0.325$ |
| Phase angle (h) | 1.32 ± 0.31 | 1.91 ± 0.16 | 1.48 ± 0.25 | $F(2,34) = 1.46$, $p = 0.246$ |
| DLMOffset (h) | 8.35 ± 0.45 | 9.04 ± 0.26 | 8.69 ± 0.22 | $F(2,35) = 1.32$, $p = 0.280$ |
| Amplitude (pg/ml) | 11.00 ± 1.44 | 10.71 ± 1.02 | 9.92 ± 1.32 | $F(2,34) = 0.57$, $p = 0.573$ |
| AUC (ng-h/L) | 3.17 ± 0.36 | 3.78 ± 0.41 | 2.96 ± 0.39 | $F(2,34) = 1.77$, $p = 0.185$ |

DLMOnset = dim light melatonin onset; DLMOffset = dim light melatonin offset; AUC = area under the curve. Values represent mean ± standard errors for each condition.

For a more detailed inspection, the individual values of the DLMOnset, DLMOffset, and the amplitude on day ten of treatment are presented in Figure S2.

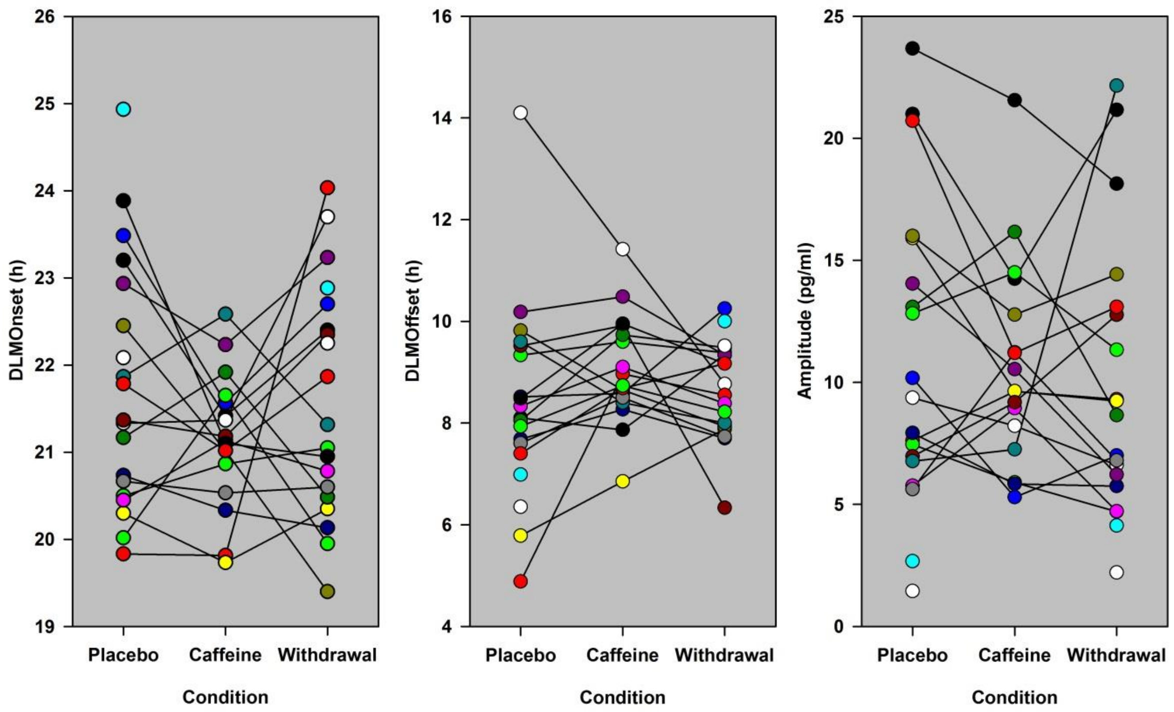


**Figure S2.** Depicted are the individual values for the melatonin parameters DLMOnset (left), DLMOffset (center), and amplitude (right) assessed on day ten of treatment. The exclusion of a visually identified extreme value in the analyses of DLMOffset (depicted as white point in the figure) reveals a significant delay in the caffeine condition compared to placebo (main effect of condition: $F_{2,35} = 3.75$; $p = 0.034$; post-hoc tests: $p = 0.030$).

### PSG recordings

For recordings, we used a portable V-Amp device (Brain Products GmbH, Gilching, Germany) and Grass Gold Cup Electrodes. Applied according to the standard international 10-20 system, two electro-oculargraphic, two electro-myographic, two electro-cardiographic signals were recorded, together with six derivations from the frontal, central, and occipital regions (F3, F4, C3, C4, O1, O2) referenced against the linked mastoids (A1, A2). Data were recorded with a sampling rate of 500 Hz and filtered online by applying a notch filter at 50 Hz. Epochs of 30 seconds were visually scored. Additionally, one third of the data have been scored by a second trained staff member in order to ensure a continuous scoring agreement of at least 85%.

Salivary caffeine levels

As expected, caffeine levels were higher in the caffeine condition compared to both the withdrawal and placebo condition (main effect of condition: $F_{2,121} = 185.16$; $p < 0.0001$; post-hoc tests: $p < 0.0001$) while levels during withdrawal were increased compared to the placebo condition (post-hoc tests: $p < 0.035$; mean ± SD: placebo: 18.52 ± 84.80 ng/ml; caffeine: 3024.18 ± 2163.68 ng/ml; withdrawal: 296.81 ± 683.42 ng/ml). As illustrated in Figure S3, this general pattern was modulated by time (interaction condition x time ($F_{22,237}$ = 2.76; $p < 0.0001$)).

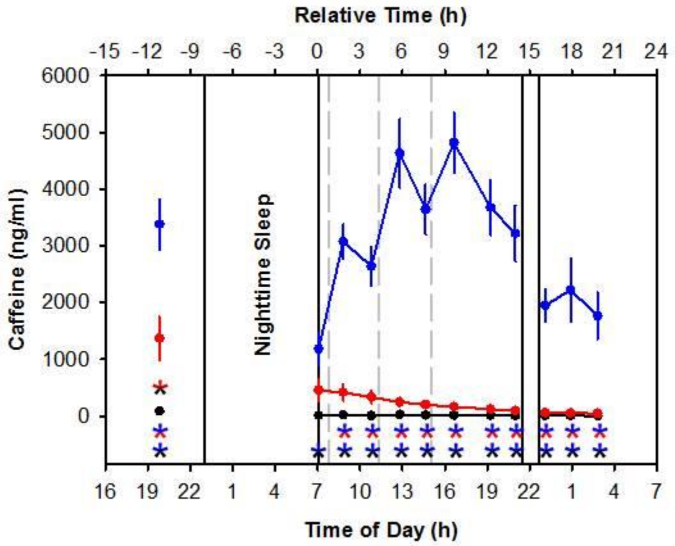


**Figure S3.** Time course of caffeine levels across the 43 h under controlled laboratory conditions. Pill administrations (placebo or caffeine) are indicated with dashed lines. Asterisks indicate significant ($p < 0.05$) post-hoc comparisons of the interaction effect condition x time, corrected according to [5] for multiple comparisons. Color-coding of the asterisks indicates the differing conditions (black: placebo, blue: caffeine, red: withdrawal).
